# Genomic data reveal similar genetic differentiation between species with vastly different dispersal capabilities and life histories

**DOI:** 10.1101/044073

**Authors:** Steve Jordan, Brian K. Hand, Scott Hotaling, Amanda DelVecchia, Rachel Malison, Clark Nissley, Jack Stanford, Gordon Luikart

## Abstract

Little is known about the life histories, population connectivity, or dispersal mechanisms of shallow groundwater organisms. Here we used RAD-seq to analyze population structure in two aquifer species: *Paraperla frontalis*, a stonefly with groundwater larvae and aerial adults, and *Stygobromus* sp., a groundwater-obligate amphipod. We found similar levels of connectivity in each species between floodplains separated by ~70 river km in the Flathead River basin of NW Montana, USA. Given that *Stygobromous* lacks the aboveground life stage of *P. frontalis*, our findings suggest that aquifer-obligate species might have previously unrecognized dispersal capacity.

## 1. Introduction

In 1974, Stanford and Gaufin [1] reported stoneflies in an alluvial aquifer supplying domestic water to a Montana community. Researchers have since documented diverse communities of macroinvertebrates, meiofauna, and microbes in shallow aquifers worldwide, and explored the conservation and ecological importance of floodplain habitats [e.g., 2,3]. These communities include insects that spend time above ground (amphibionts), as well as taxa (e.g. crustaceans, oligochaetes, and mites) that never leave interstitial spaces in aquifers (stygobionts). These animals occur up to 10 meters beneath riverine floodplains and as many as two km from the main river channels [4]. Amphibiotic stoneflies spend 1-3 years maturing in the aquifer before emerging as adults with an aerial lifespan of only a few days [5]. The life cycles of many stygobionts, including *Stygobromus*, are largely unknown.

Shallow aquifers offer many challenges to resident organisms including geologically bounded isolation, no light, variable water flow, and reduced availability of carbon, other nutrients, and oxygen [6,7]. Recent stygobiont research has focused on the ecology of shallow groundwater environments, noting the variable influence of many factors on spatial distribution, including bedrock geology, soil permeability, water chemistry and quality, groundwater levels, adjacent surface flows, riparian vegetation, and climate [8–11]. A recent study documented the role of methane in subterranean food webs [7]. Previous genetic studies identified widespread, long-term barriers to dispersal by groundwater species, even within drainages, despite potentially linking floods [12–15].

The ability of groundwater organisms to actively disperse within and between adjacent watersheds remains unclear. Life history surely plays a significant role in population connectivity, with the retention of ephemeral, winged life stages by amphibionts (e.g., stoneflies) offering dispersal advantages over taxa that never leave the groundwater (e.g., amphipods). Current genomic tools facilitate the study of local adaptation to atypical environments [16] and can resolve fine-scale differentiation in aquatic insects [17–19]. A better understanding of dispersal in these systems would greatly benefit biological conceptualization of connectivity along the river corridor, a major theme in river ecology and management [e.g., 20].

Here we address these issues with RAD-seq datasets for two groundwater species from two well-characterized floodplains of the Flathead River in northwestern Montana. This study design is novel and powerful, using cutting-edge techniques to compare population connectivity in two co-occurring, but ecologically distinct, species.

## Methods

### Study sites

Our samples are from two Flathead River floodplains that are separated by ~70 km: the Nyack and the Kalispell (Fig. 1). Both locations are underlain by high-porosity alluvial aquifers that are entirely recharged by river water. The aquifers are likely not connected in the subsurface because the floodplains are bounded by bedrock knickpoints [21], and each is known to contain a diverse array of meiofauna and macroinvertebrates [7,22]. These floodplains have been the focus of long-term research [e.g., 23], and permanent wells allow sampling of groundwater habitats.

**Figure 1.**
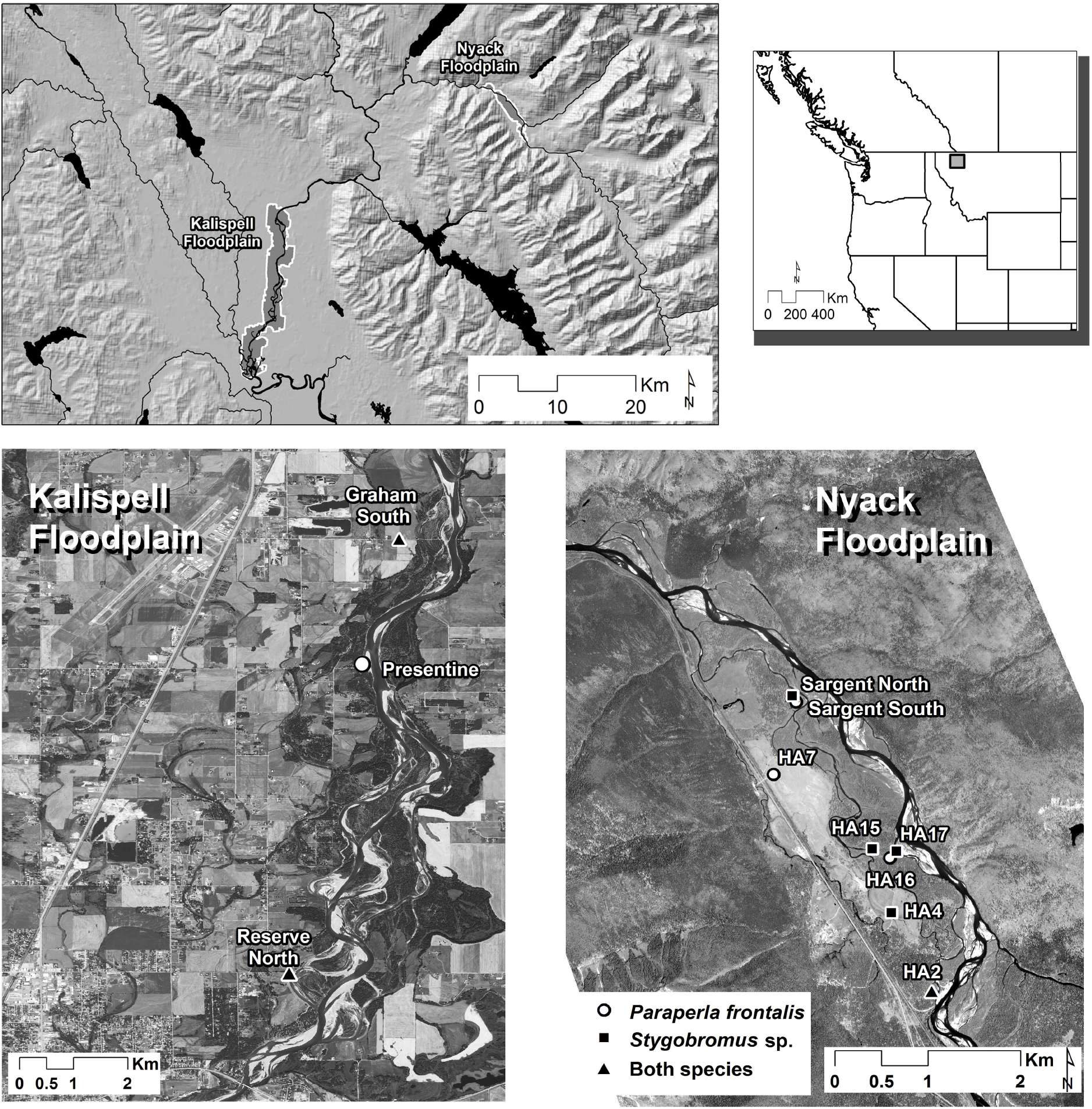
Sampling sites, from region to floodplain to individual wells, for two species of groundwater invertebrates included in this study.

### Taxa and sampling

We sampled two taxa that exemplify major life history strategies: *Paraperla frontalis*, a hyporheic stonefly with a winged adult stage, and *Stygobromus*, a blind, pigmentless, groundwater-obligate crustacean. We hypothesized that the stonefly would display higher gene flow and less genetic structure between floodplains than the groundwater amphipod.

We collected specimens from seven permanent wells in June of 2011 and 2012, though both species were not always sampled from the same wells (Fig. 1). Because of limited sampling, all *Stygobromus* from Kalispell were considered one population in analyses. We extracted DNA from 96 individuals of each species and confirmed species identifications by COI barcoding before proceeding with RAD-seq.

### SNP calling and filtering

We prepared RAD-seq libraries for both species following standard protocols using the restriction enzyme SbfI and unique 6-bp barcodes [24]. We sequenced 192 samples on two lanes of an Illumina HiSeq 2500 with 100-bp, single-end chemistry. Raw sequences were quality filtered with 90% of the bases required to have a quality score ≥20. Reads for *Stygobromus* were further trimmed to 80 base pairs to maximize read quality. We used the process_radtags script in Stacks v.1.19 [25] to demultiplex reads by barcode, removing any with an uncalled base. We called SNPs in the program Stacks [25] with a read depth (-m) of 5 and a maximum of two mismatches (-n) per locus and between catalog loci (-N). Additional SNP calling details are provided in Supplementary Materials. When necessary, we used PGDSpider 2.1.1.1 [26] for data conversion. To investigate the influence of data scale (SNP number) and missing data on our results, we constructed four datasets for downstream analyses. All filtering steps applied to both species and all datasets received the baseline filtering described above. (1) We removed all individuals with >50% missing data, and all loci genotyped in < 60% of individuals. We also removed all loci that were not present in at least 50% of individuals in each population. (2) No additional filters. (3) We removed loci genotyped in <25% of individuals. (4) We removed loci genotyped in <75% of individuals. For datasets 2-4, we also removed any individuals with more than one standard deviation of missing data above the mean in the raw SNP dataset.

### Pair-wise differentiation and population structure

We calculated pairwise F_ST_ values using GENEPOP [27,28] for all population pairs using dataset #1. Next, we explored population structure in two ways for four datasets. We used Structure v2.3.4 [29], a Bayesian iterative algorithm, and a discriminant analysis of principal components (DAPC) implemented in the R package *adegenet* [30]. For both methods, we tested *K*=1-7 for *P. frontalis* and *K*=1-6 for *Stygobromus*. Structure analyses were performed on dataset #1 and DAPC analyses were performed on datasets #2-4. Complete details of population structure analyses are provided in Supplementary Materials.

## Results

### Sequencing and genotyping

We observed lower coverage depth and fewer SNPs despite a much larger SNP catalog (e.g., 2-3x) for *Stygobromus* versus *P. frontalis*. This is likely because the genome of *Stygobromus* is much larger than *P. frontalis* [see 31]. For each dataset, we identified: (1) 806 SNPs for 90 *P. frontalis* and 314 SNPs for 50 *Stygobromus*, (2) 3,187 SNPs for *P. frontalis* and 543 SNPs for *Stygobromus*, (3) 1,175 SNPs for *P. frontalis* and 476 SNPs *Stygobromus*, and (4) 279 SNPs for *P. frontalis* and 167 SNPs for *Stygobromus*. For datasets 2-4, we included 84 and 46 individuals of *P. frontalis* and *Stygobromus*, respectively (Table 1).

**Table 1.** Sample sizes of the amphipod *Stygobromus* sp. and the stonefly *Paraperla frontalis*, including total numbers of individuals with DNA initially extracted and those that met the quality requirements for the final RAD-seq data sets (#1 and #2-4).

| Floodplain | Population | *Stygobromus* sp. |  |  | *P. frontalis* |  |  |
| --- | --- | --- | --- | --- | --- | --- | --- |
|  |  | Extracted | #1 | #2-4 | Extracted | #1 | #2-4 |
| Nyack | Sargent South |  |  |  | 26 | 25 | 22 |
|  | Sargent North | 15 | 3 | 3 |  |  |  |
|  | Ha2 | 14 | 15 | 15 | 14 | 12 | 12 |
|  | Ha4 | 13 | 3 | 2 |  |  |  |
|  | Ha7 |  |  |  | 14 | 14 | 14 |
|  | Ha15 | 18 | 11 | 9 |  |  |  |
|  | Ha16 |  |  |  | 14 | 13 | 10 |
|  | Ha17 | 8 | 8 | 8 |  |  |  |
| Kalispell | Graham South | 16 | 1 | 1 | 14 | 12 | 12 |
|  | Reserve North | 12 | 9 | 8 | 8 | 8 | 8 |
|  | Presentine Bar |  |  |  | 6 | 6 | 6 |
| Total |  | 96 | 50 | 46 | 96 | 90 | 84 |

### Pair-wise differentiation and population structure

We calculated pairwise F_ST_ values within and between floodplains for both species using dataset 1 (Table 2). However, because we kept just 10 *Stygobromus* individuals from the two Kalispell wells, we did not calculate pairwise F_ST_ within that floodplain. We found genetic differentiation between floodplains to be low for both species (F_ST_=0.004 for *Paraperla* and F_ST_=0.000 for *Stygobromus*). These low F_ST_ were likewise reflected in low population pairwise F_ST_ values (Table 2).

**Table 2.**
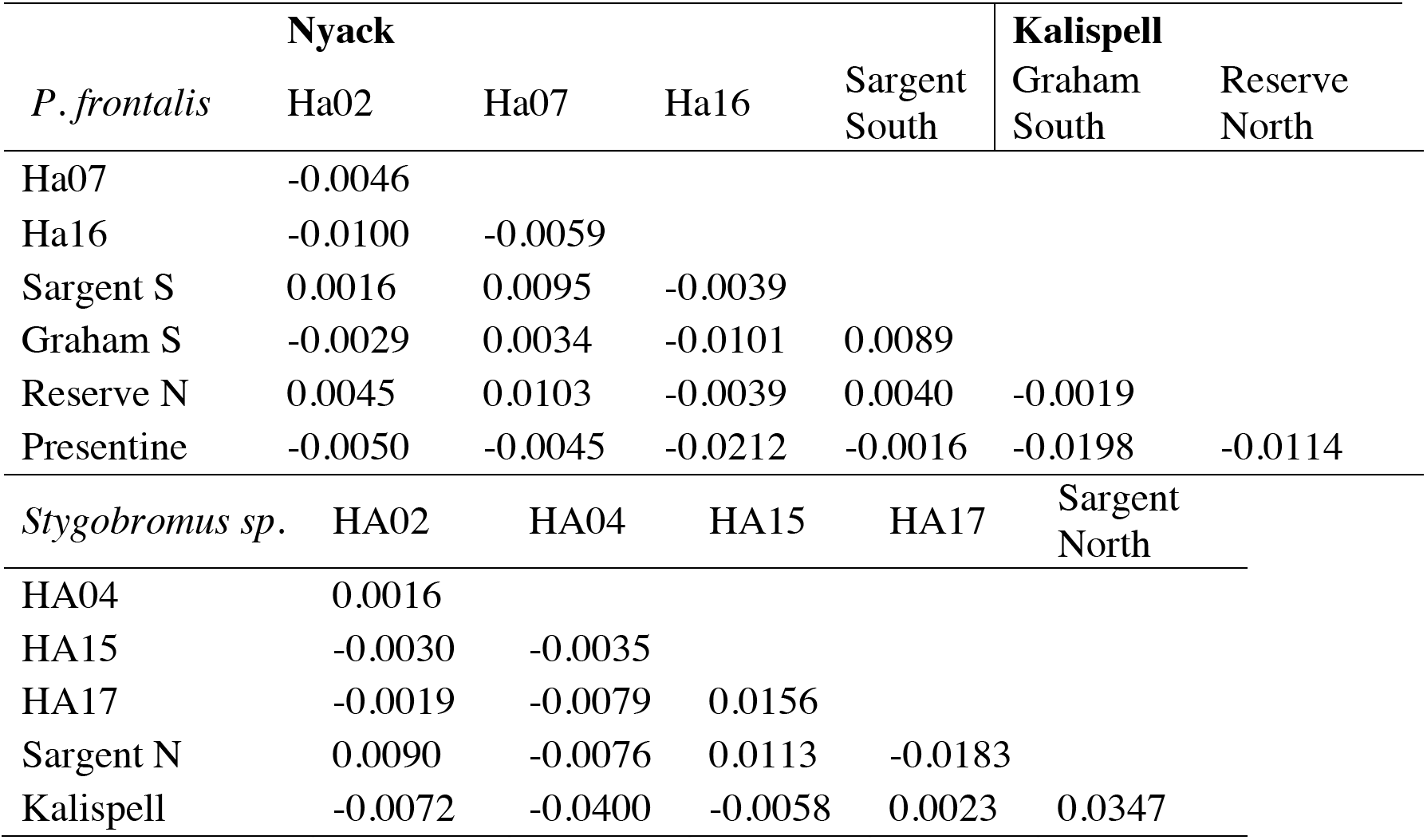
Pairwise *F*_ST_ values for samples of *Paraperla frontalis* and *Stygobromus* sp. as calculated from data set #1 which included 806 SNPs for 90 *P. frontalis* individuals and 314 SNPs for 50 *Stygobromus* sp. individuals. Bold names indicate different floodplains.

|  | **Nyack** |  |  |  | **Kalispell** |  |
| --- | --- | --- | --- | --- | --- | --- |
| *P. frontalis* | Ha02 | Ha07 | Ha16 | Sargent South | Graham South | Reserve North |
| Ha07 | -0.0046 |  |  |  |  |  |
| Ha16 | -0.0100 | -0.0059 |  |  |  |  |
| Sargent S | 0.0016 | 0.0095 | -0.0039 |  |  |  |
| Graham S | -0.0029 | 0.0034 | -0.0101 | 0.0089 |  |  |
| Reserve N | 0.0045 | 0.0103 | -0.0039 | 0.0040 | -0.0019 |  |
| Presentine | -0.0050 | -0.0045 | -0.0212 | -0.0016 | -0.0198 | -0.0114 |
| *Stygobromus* sp. | HA02 | HA04 | HA15 | HA17 | Sargent North |  |
| HA04 | 0.0016 |  |  |  |  |  |
| HA15 | -0.0030 | -0.0035 |  |  |  |  |
| HA17 | -0.0019 | -0.0079 | 0.0156 |  |  |  |
| Sargent N | 0.0090 | -0.0076 | 0.0113 | -0.0183 |  |  |
| Kalispell | -0.0072 | -0.0400 | -0.0058 | 0.0023 | 0.0347 |  |

For population structure analyses, we identified optimal values of *K*=2 for *P. frontalis* and *K*=5 for *Stygobromus* (Figs. S1–2). DAPC analyses revealed similar levels of population structure for both species. For *P. frontalis*, the optimal *K* ranged from *K*=2 (datasets 2-3) to *K*=3 (dataset 4; Fig. 2A). For *Stygobromus*, the best-fit DAPC *K* was either *K*=5 (dataset 2), *K*=4 (dataset 3), or *K*=3 (dataset 4; Fig. 2B), with all *K*’s highlighting no obvious patterns of geographic structuring.

**Figure 2.**
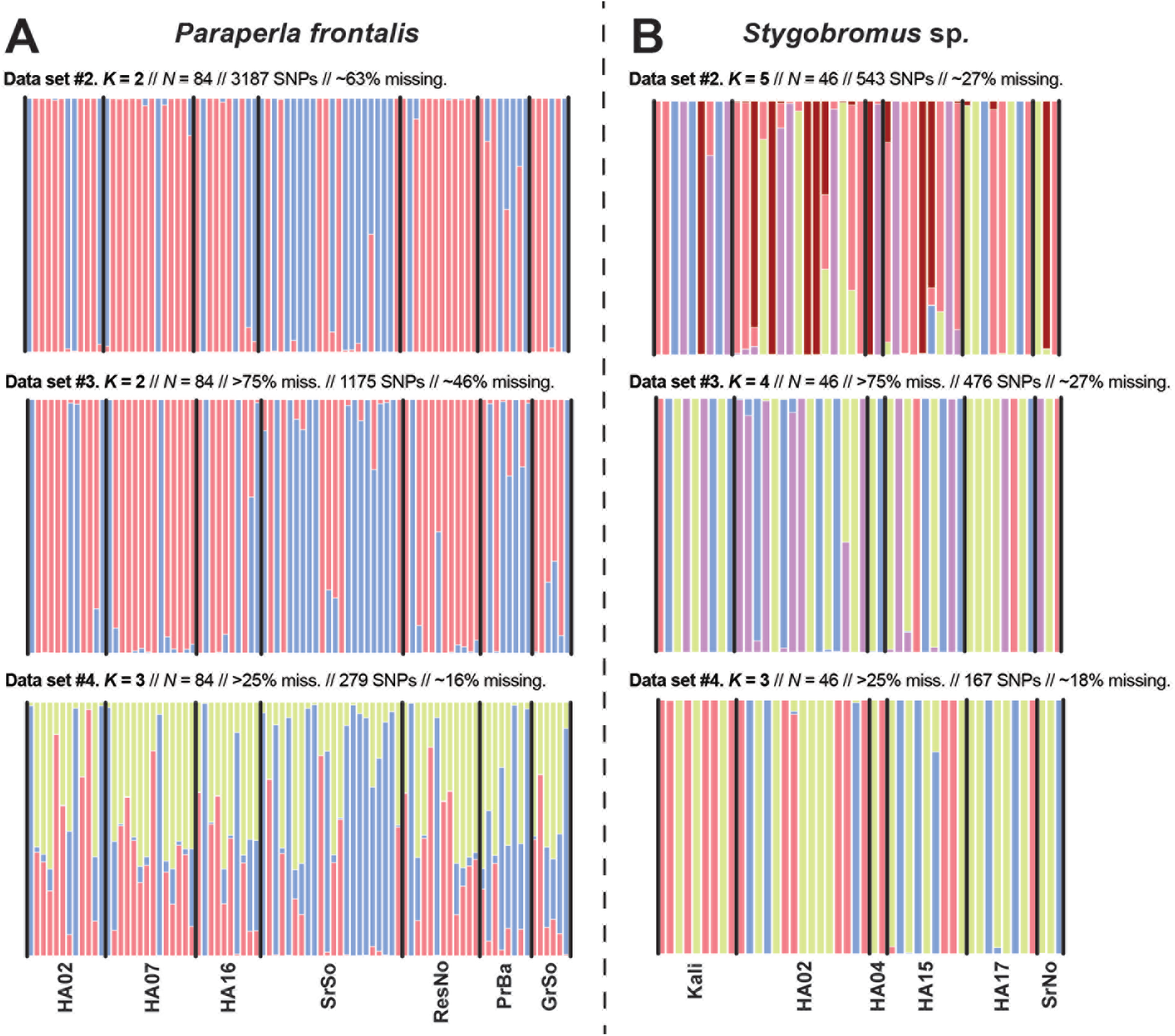
Comparisons of population structure for DAPC analyses of data sets #2-4 which include varying numbers of SNPs for A) *Paraperla frontalis* and B) *Stygobromus sp*. For each plot, the number of clusters supported (*K*), filters employed, number of SNPs, and percent of missing data are included. Filter abbreviations include: >75% miss. and >25% miss. = removal of any loci with greater than 75% or 25% missing data, respectively. Each vertical bar represents one individual, and best-fit *K*’s were identified by plotting the Bayesian Information Criterion (BIC) for A) *K* = 1-7 and B) *K* = 1-6. Lower BICs indicate higher model support.

## Discussion

Shallow alluvial aquifers are refugia for diverse, functionally unique species that are inherently difficult to study. Thus, we are only beginning to understand such fundamental attributes as their dispersal ability, population structure, mating habits, and adaptability. Our study compared winged stoneflies with groundwater amphipods, species we expected to exhibit vast differences in their dispersal capacity. Using hundreds to thousands of loci, we have shown a lack of regional structure in both our study species, a surprising result given their different life histories, predicted dispersal differences, and the barriers (geologic and geographic) separating populations.

There are several possible explanations for this result. First, the deep bedrock, phreatic, water channels underlying this region may connect distant floodplains, thus facilitating dispersal of one or both of our taxa. However, entry of groundwater taxa into such channels may be limited because they lack the direct surface-to-groundwater connection present in the hyporheic zone. Furthermore, the environment of these channels would be challenging to shallow aquifer taxa, with longer residence times and potentially limiting temperatures, carbon availability, and oxygen levels. Second, it is possible that *Stygobromus* individuals irregularly enter the river current through upwelling and passively disperse beyond bedrock nickpoints to become established in downstream floodplains. Flood events may contribute to their movement as well, though they would still have to be brought out of the hyporheos in some way. Even very low dispersal rates could lead to genetic homogeneity [32].

In summary, we applied powerful genomic tools to a fundamental biological question in a long-studied system and found a novel, surprising result. Our results clearly indicate that differences in dispersal capacity cannot be definitively assumed from life history differences, even something as extreme as purely subterranean versus winged life stages. Ongoing work will extend these results with more robust taxonomic, genomic, and geographic sampling. In the future, we expect to better resolve the historical and ecological drivers of groundwater genomic and phylogenetic diversity.

## Acknowledgements

We thank all field and lab technicians, as well as Mike Miller, Sean O’Rourke, Omar Ali, and Laney Hayssen for laboratory assistance.

## Author contributions

SJ, JS, and GL designed the study. SJ, CN, JS, and GL did field and/or lab work. SJ, BKH, and SH performed analyses. All authors contributed to manuscript preparation.

## Data accessibility

Data will be deposited in GenBank and other repositories.

## Ethical statement

Invertebrates were sampled humanely under permits from USGS, FWS, and the state of Montana.

---

**Figure S1.**
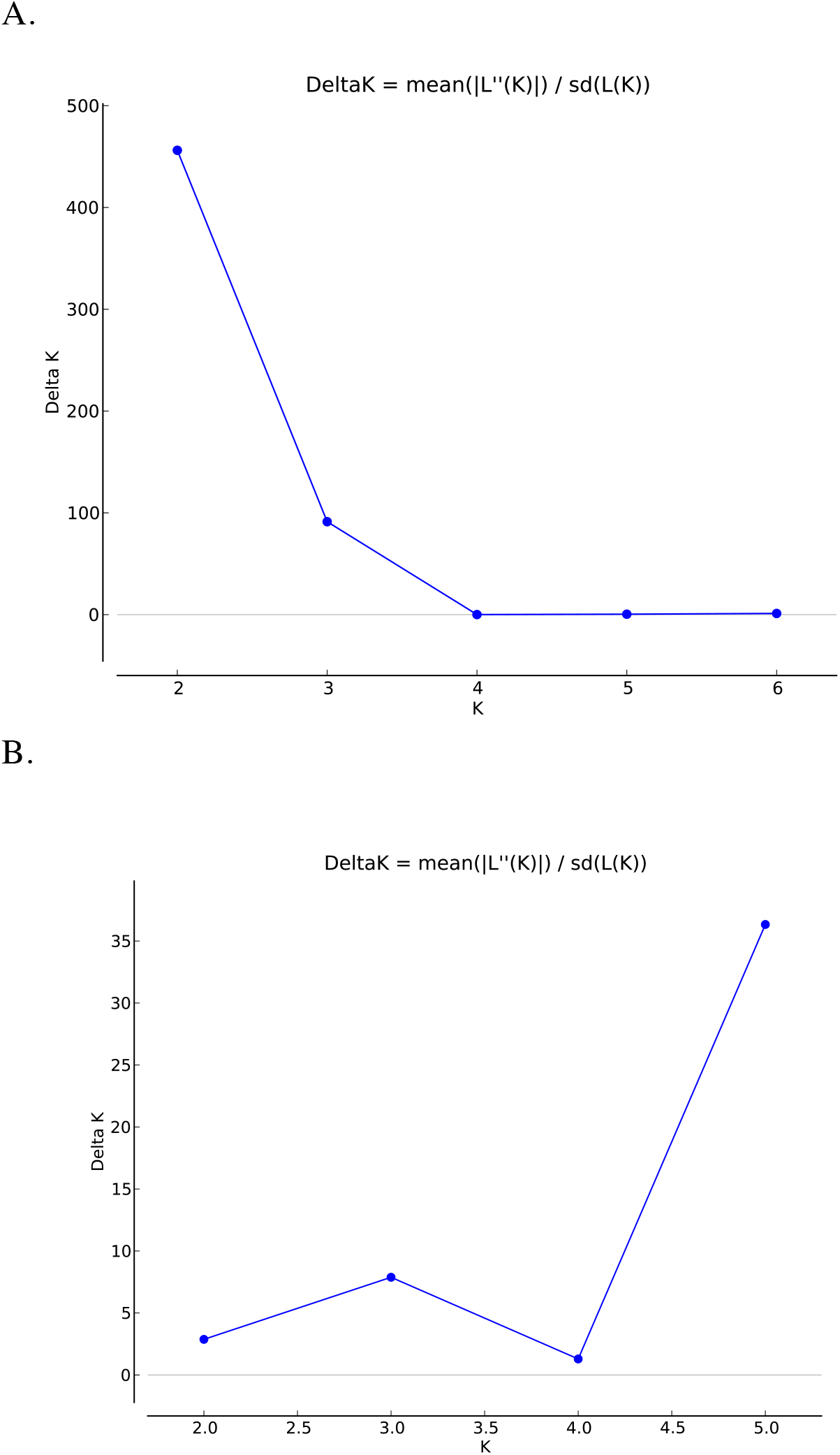
Results of Structure Harvester analysis showing optimal values of K for A) *Paraperla frontalis* and B) *Stygobromus* sp.

**Figure S2.**
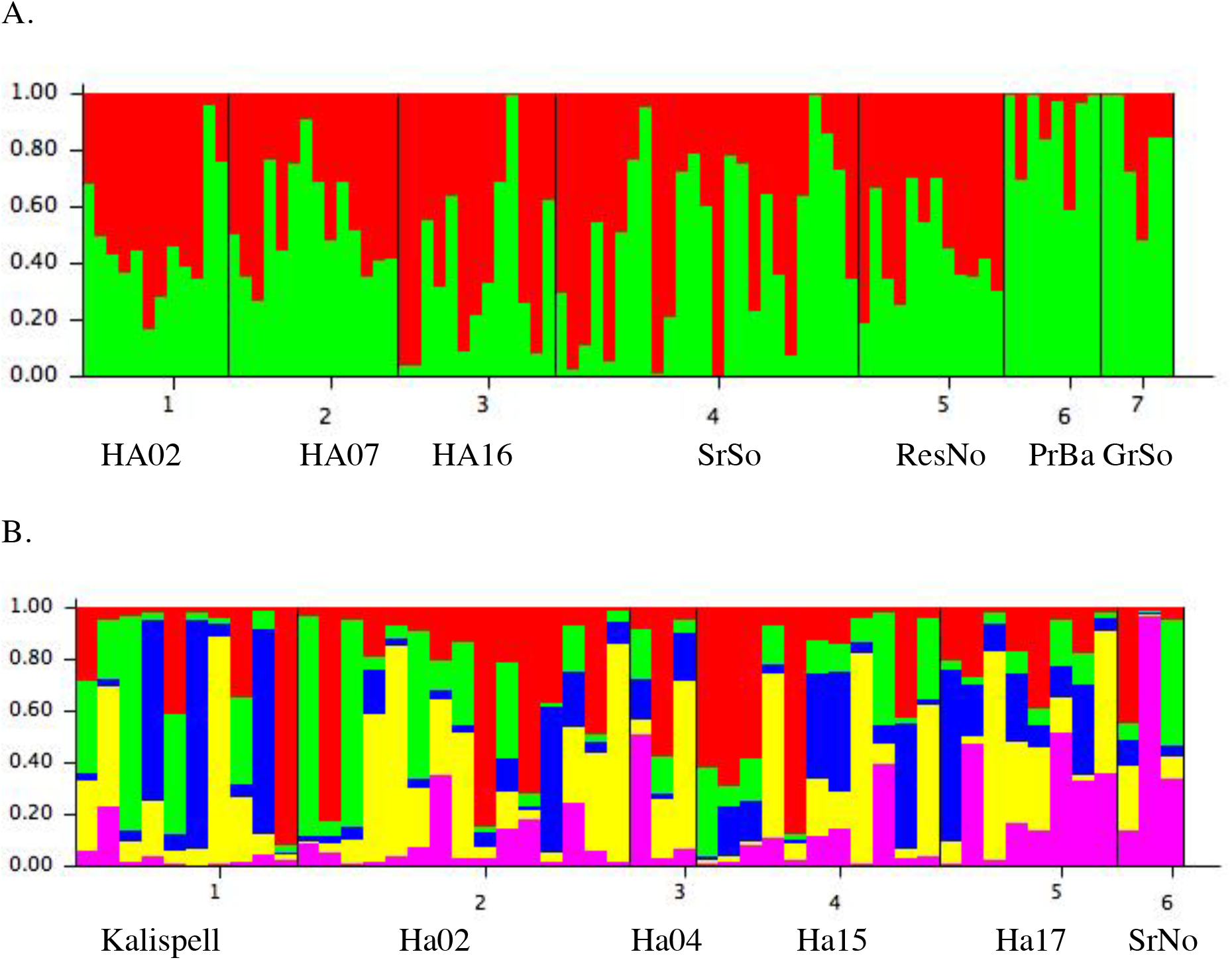
Results of Structure analysis for A) *Paraperla frontalis* and B) *Stygobromus* sp., with K=2 and K=5, respectively.

## References

1. Stanford JA, Gaufin AR. 1974 Hyporheic communities of two montana rivers. Science 185, 700–702. (doi:10.1126/science.185.4152.700)

2. Boulton AJ, Datry T, Kasahara T, Mutz M, Stanford JA. 2010 Ecology and management of the hyporheic zone: stream–groundwater interactions of running waters and their floodplains. J. North Am. Benthol. Soc. 29, 26–40. (doi:10.1899/08-017.1)

3. Hauer FR, Locke H, Dreitz VJ, Hebblewhite M, Lowe WH, Muhlfeld CC, Nelson CR, Proctor MF, Rood SB. 2016 Gravel-bed river floodplains are the ecological nexus of glaciated mountain landscapes. Sci. Adv. 2, e1600026–e1600026. (doi:10.1126/sciadv.1600026)

4. Stanford JA, Ward J V. 1988 The hyporheic habitat of river ecosystems. Nature 335, 64–66. (doi:10.1038/335064a0)

5. Stewart KW, Stark BP. 2002 Nymphs of North American Stonefly Genera (Plecoptera). Second Edition. Stewart, Kenneth W.; University of North Texas, Denton, USA, USA.: Caddis Press.

6. Tockner K, Pusch M, Borchardt D, Lorang MS. 2010 Multiple stressors in coupled river-floodplain ecosystems. Freshw. Biol. 55, 135–151. (doi:10.1111/j.1365-2427.2009.02371.x)

7. DelVecchia AG, Stanford JA, Xu X. 2016 Ancient and methane-derived carbon subsidizes contemporary food webs. Nat. Commun. 7, 13163. (doi:10.1038/ncomms13163)

8. Larned T. S, Unwin MJ, Boustead NC. 2015 Ecological dynamics in the riverine aquifers of a gaining and losing river. Freshw. Sci. 34, 245–262. (doi:10.1086/678350)

9. Korbel KL, Hose GC. 2015 Habitat, water quality, seasonality, or site? Identifying environmental correlates of the distribution of groundwater biota. Freshw. Sci. 34, 329–343. (doi:10.1086/680038)

10. Stubbington R, Boulton AJ, Little S, Wood PJ. 2015 Changes in invertebrate assemblage composition in benthic and hyporheic zones during a severe supraseasonal drought. Freshw. Sci. 34, 344–354. (doi:10.1086/679467)

11. Johns T, Jones JI, Knight L, Maurice L, Wood P, Robertson A. 2015 Regional-scale drivers of groundwater faunal distributions. Freshw. Sci. 34, 316–328. (doi:10.1086/678460)

12. Finston TL, Johnson MS, Humphreys WF, Eberhard SM, Halse SA. 2007 Cryptic speciation in two widespread subterranean amphipod genera reflects historical drainage patterns in an ancient landscape. Mol. Ecol. 16, 355–365. (doi:10.1111/j.1365- 294X.2006.03123.x)

13. Lefebure T, Douady CJ, Gouy M, Trontelj P, Briolay J, Gibert J. 2006 Phylogeography of a subterranean amphipod reveals cryptic diversity and dynamic evolution in extreme environments. Mol. Ecol. 15, 1797–1806. (doi:10.1111/j.1365-294X.2006.02888.x)

14. Cooper SJB, Bradbury JH, Saint KM, Leys R, Austin AD, Humphreys WF. 2007 Subterranean archipelago in the Australian arid zone: mitochondrial DNA phylogeography of amphipods from central Western Australia. Mol. Ecol. 16, 1533–1544. (doi:10.1111/j.1365-294X.2007.03261.x)

15. Cooper SJB, Saint KM, Taiti S, Austin AD, Humphreys WF. 2008 Subterranean archipelago: mitochondrial DNA phylogeography of stygobitic isopods (Oniscidea: Haloniscus) from the Yilgarn region of Western Australia. Invertebr. Syst. 22, 195–203. (doi:10.1071/IS07039)

16. Luikart G, England PR, Tallmon D, Jordan S, Taberlet P. 2003 The power and promise of population genomics: from genotyping to genome typing. Nat. Rev. Genet. 4, 981–994.

17. Boumans L, Hogner S, Brittain J, Johnsen A. 2017 Ecological speciation by temporal isolation in a population of the stonefly Leuctra hippopus (Plecoptera, Leuctridae). Ecol. Evol. 7, 1635–1649. (doi:10.1002/ece3.2638)

18. Hotaling S, Muhlfeld CC, Giersch JJ, Ali OA, Jordan S, Miller MR, Luikart G, Weisrock DW. 2018 Demographic modelling reveals a history of divergence with gene flow for a glacially tied stonefly in a changing post-Pleistocene landscape. J. Biogeogr. (doi:10.1111/jbi.13125)

19. Dussex N, Chuah A, Waters JM. 2016 Genome-wide SNPs reveal fine-scale differentiation among wingless alpine stonefly populations and introgression between winged and wingless forms. Evolution (N. Y). 70, 38–47. (doi:10.1111/evo.12826)

20. Stanford JA, Lorang MS, Hauer FR. 2005 The shifting habitat mosaic of river ecosystems. Travaux. Assoc. Int. Limnol. Theor. Appl. 29, 1–14.

21. Hauer Fr, Stanford A. J, Lorang S. M. 2007 Pattern and process in Northern Rocky Mountain headwaters: ecological linkages in the headwaters of the Crown of the Continent. J. Am. Water Resour. Assoc. 43, 104–117.

22. Stanford JA, Ward J V, Ellis BK. 1994 Ecology of the alluvial aquifers of the Flathead River, Montana. In Groundwater Ecology (ed JD Gibert Dan L. Stanford,Jack, A.), pp. 367–390. San Diego, California: Academic Press.

23. Helton M. A, Poole C. G, Payn A. R, Izurieta C, Stanford A. J. 2014 Relative influences of the river channel, floodplain surface, and alluvial aquifer on simulated hydrologic residence time in a montane river floodplain. Geomorphology 205, 17–26.

24. Miller MR, Brunelli JP, Wheeler PA, Liu S, Rexroad, III CE, Palti Y, Doe CQ, Thorgaard GH. 2012 A conserved haplotype controls parallel adaptation in geographically distant salmonid populations. Mol. Ecol. 21, 237–249. (doi:10.1111/j.1365-294X.2011.05305.x)

25. Catchen J, Hohenlohe PA, Bassham S, Amores A, Cresko WA. 2013 Stacks: an analysis tool set for population genomics. Mol. Ecol. 22, 3124–3140. (doi:10.1111/mec.12354)

26. Lischer HEL, Excoffier L. 2012 PGDSpider: An automated data conversion tool for connecting population genetics and genomics programs. Bioinformatics 28, 298–299. (doi:10.1093/bioinformatics/btr642)

27. Raymond M, Rousset F. 1995 GENEPOP (version 1.2): population genetics software for exact tests and ecumenicism. J. Hered. 86, 248–249.

28. Rousset F. 2008 GENEPOP ‘007: a complete re-implementation of the GENEPOP software for Windows and Linux. Mol. Ecol. Resour. 8, 103–106.

29. Pritchard K. J, Stephens M, Donnelly P. 2000 Inference of population structure using multilocus genotype data. Genetics 155, 945–959.

30. Jombart T. 2008 adegenet: a R package for the multivariate analysis of genetic markers. Bioinformatics 24, 1403–1405. (doi:10.1093/bioinformatics/btn129)

31. Rees DJ, Dufresne F, Glemet H, Belzile C. 2007 Amphipod genome sizes: first estimates for Arctic species reveal genomic giants. Genome 50, 151–158. (doi:10.1139/G06-155)

32. Allendorf FW, Luikart G. 2007 Conservation and the Genetics of Populations. Malden, MA: Wiley-Blackwell.

